# High-throughput single-cell isolation of *Bifidobacterium* strains from the gut microbiome

**DOI:** 10.1101/2025.07.23.666462

**Authors:** Lam Hai Ha, Yue Yuan On, Clarice Pohan, Jungwon Lee, Shaun Hong Chuen How, Yik-Ying Teo, Henning Seedorf, Jean-Sebastien Gounot, Niranjan Nagarajan

**Affiliations:** Genome Institute of Singapore (GIS), Agency for Science, Technology and Research (A*STAR), 60 Biopolis Street, Genome, Singapore, 138672, Republic of Singapore; Life Sciences Institute, National University of Singapore, Singapore, 117456, Singapore; Saw Swee Hock School of Public Health, National University of Singapore, 12 Science Drive 2, Singapore, 117549, Singapore; Department of Statistics and Applied Probability, National University of Singapore, Singapore, 117546, Singapore; Temasek Lifesciences Laboratory, 1 Research Link, Singapore, 117604, Singapore; Department of Biological Sciences, National University of Singapore, Singapore, 117558, Singapore; Department of Biochemistry, Yong Loo Lin School of Medicine, National University of Singapore, Singapore

**Keywords:** high-throughput isolation, culturomics, *Bifidobacterium*

## Abstract

While metagenomic studies can highlight strain-level diversity within microbial communities, the diversity obtained is often incomplete. Moreover, their utility for phenotypic characterizations remains hampered without the subjacent, systematic isolation procedures required with traditional culturomics. In this work, we examined the capabilities of a commercially available high-throughput single-cell dispensing solution to selectively target and isolate diverse strains of a genus of interest, *Bifidobacterium*, from fecal samples. The general performance of the single-cell dispenser was first assessed, revealing a low doublet frequency of 11.5% and an ability to preserve global genus diversity when a mixed culture of *Bifidobacterium* was dispensed. Culturing-related factors including the use of an effective selection medium, such as the Bifidus Selective Medium supplemented with mupirocin (BSM-MUP), and the length of incubation were found to be critical in determining isolation success. Leveraging these results, we obtained a total of 622 viable isolates from five Singaporean fecal samples, among which ∼98.7% were bifidobacteria. Whole-genome sequencing of 96 isolates revealed six different *Bifidobacterium* species with both inter- and intra-subject lineage diversity, and the majority of the assemblies were not previously captured using metagenomic sequencing. Our findings validate the ability of high-throughput culturomics to recover diverse, novel bacterial strains and open up the possibility to robustly interrogate their functional characteristics, advancing our understanding of important microbiomes.

**IMPORTANCE:** The field of microbial culturomics is still in its early stages. Enhancing our ability to isolate and phenotypically test bacterial strains from their multicellular environment is crucial for advancing microbiome research and healthcare development. Given the time- and cost-inefficiencies of traditional culturing methods, a more efficient, high-throughput approach to obtain isolates is needed.

In the present study, we assessed a single-cell dispensing platform and developed a workflow to isolate diverse *Bifidobacterium* strains from fecal samples. We demonstrated here the capability of this novel technology to efficiently obtain hundreds of isolates of a targeted organism, covering both species and strain diversities. This generalizable and scalable method allows for the high-throughput recovery of microbes with little optimization needed for novel targets, providing a fundamental step in improving the culturomic framework to complement metagenomic approaches and enable isolate-level functional studies of important microbiomes.

## INTRODUCTION

Culturomics is an essential tool for understanding the diversity of microbial communities (1, 2). By leveraging different culturing media and conditions, it has been successfully used to cultivate bacteria in several environments, such as plant, soil (3–6) and human microbiota (7–10), identifying previously unknown bacteria and thereby filling the microbial “dark matter” gap in metagenomic approaches (7, 11–15). Another advantage of culturomics is the better characterization of bacteria both phenotypically and genotypically. Culturomics yields live isolates that enable direct assessment of microbial phenotypes such as metabolic profiles, antibiotic resistance, or host environment interactions that cannot be fully accessed through sequencing alone (16–19). Additionally, the ability to deeply sequence individual pure isolates results in more accurate and complete genome assemblies, especially for less abundant microbes (20–22). This high-resolution genomic data would allow for the discovery of novel species and strain-level variations, including person- and population-specific signatures.

Despite its utility, the application of culturomics remains limited by time-consuming, labour-intensive, and low-throughput bacterial isolation methods (23, 24). Thus, to increase the recovery efficiency of culturomics, automated single-cell isolation technologies have been progressively developed. Notably, microfluidic and fluorescence-activated cell sorting (FACS) systems were among the first to be used. While microfluidic systems are more affordable and physically compact, they lack control over the end products, resulting in large numbers of empty droplets (25, 26). On the other hand, FACS systems can achieve high single-cell recovery rates, but demand expertise and are more costly (27, 28). In addition, FACS systems require a large space, which makes them incompatible with enclosed environments such as the anaerobic chamber. Recently, next-generation single-cell dispensers have offered an alternative solution that balances the advantages of the earlier systems. B.SIGHT single-cell dispenser (Cytena GmBH, Germany), for instance, is a compact and user-friendly machine that utilizes an in-built camera to detect single-cell droplets for dispensing, gaining attention for bacterial isolations (23, 26, 29).

In this study, we first characterized the performance of the dispenser in terms of doublet frequency, validating its utility as a single-cell isolation platform. We then applied the single-cell dispenser in a culturomics workflow to efficiently isolate *Bifidobacterium* from fecal samples. We chose *Bifidobacterium* as our model bacterial genus as it plays many important roles in the human gut (30– 34) and, in a recent metagenomic study from our group, various uncharacterized *Bifidobacterium* strains were found in a Singaporean cohort (35). To understand the dispenser’s behaviour when handling a mixed bacterial environment, we dispensed nine *Bifidobacterium* species at once and found that the single-cell dispenser was able to capture the global diversity of the mock community. We also demonstrated that the pre-dispensing incubation period and choice of enrichment medium play a key role in determining the isolation success and recovery rates of diverse *Bifidobacterium* species. Finally, we successfully dispensed a large collection of *Bifidobacterium* isolates from five Singaporean fecal samples, with substantial lineage diversity. This study serves as proof of concept for the application of single-cell dispensers in further culturomic experiments, paving the way to population-scale microbiota isolation studies.

## RESULTS

### Characterization of the single-cell dispenser’s performance in single-cell isolation and dispensing pure *Bifidobacterium* cultures

To validate the single-cell isolation capability of the dispenser, we dispensed a mixed culture of GFP-expressing and mApple-expressing *K. pneumoniae* into six 96-well plates and detected fluorescence signals from each individual well. A majority of non-empty wells (450/477, Supplementary Table 1) displayed a single fluorescence signal (Fig. 1a, GFP- and mApple-only) while a group of isolates (27/477, Supplementary Table 1) showed both GFP and mApple fluorescence signals (Fig. 1a, GFP + mApple). This result suggests that approximately 5.66% of the dispensing instances contained at least two cells with different fluorescence expressions (Fig. 1b). Assuming a low probability of three or more cells found in one dispensing instance and that GFP- or mApple-only doublets are as likely as GFP + mApple doublets, we can estimate a total doublet dispensing frequency of ∼11.5%. Thus, given a roughly 88.5% rate of single-cell dispenses, the dispensing platform remains a reliable tool to obtain isolates for global screening from a mixed sample, but would require additional re-streaking for the construction of a pure strain database. We also tested the reusability of one of the dispenser’s components, the dispensing cartridge, to reduce operational costs for future experiments. By introducing a post-dispensing washing procedure to the dispensing workflow, the number of cells detected in the cartridge approaches near-zero levels for samples of varied cell densities (Supplementary Fig. 1a). Specifically, after three washes, we achieved a negligible, ∼0.1% false positive rate for the medium-to high-density samples and a ∼2% rate for the low-density samples, which could be further reduced to ∼0.4% upon the 5^th^ wash (Supplementary Fig. 1b). This extra washing procedure further improves the cost efficiency and utility of the single-cell dispenser in processing large numbers of samples.

**FIG 1.**
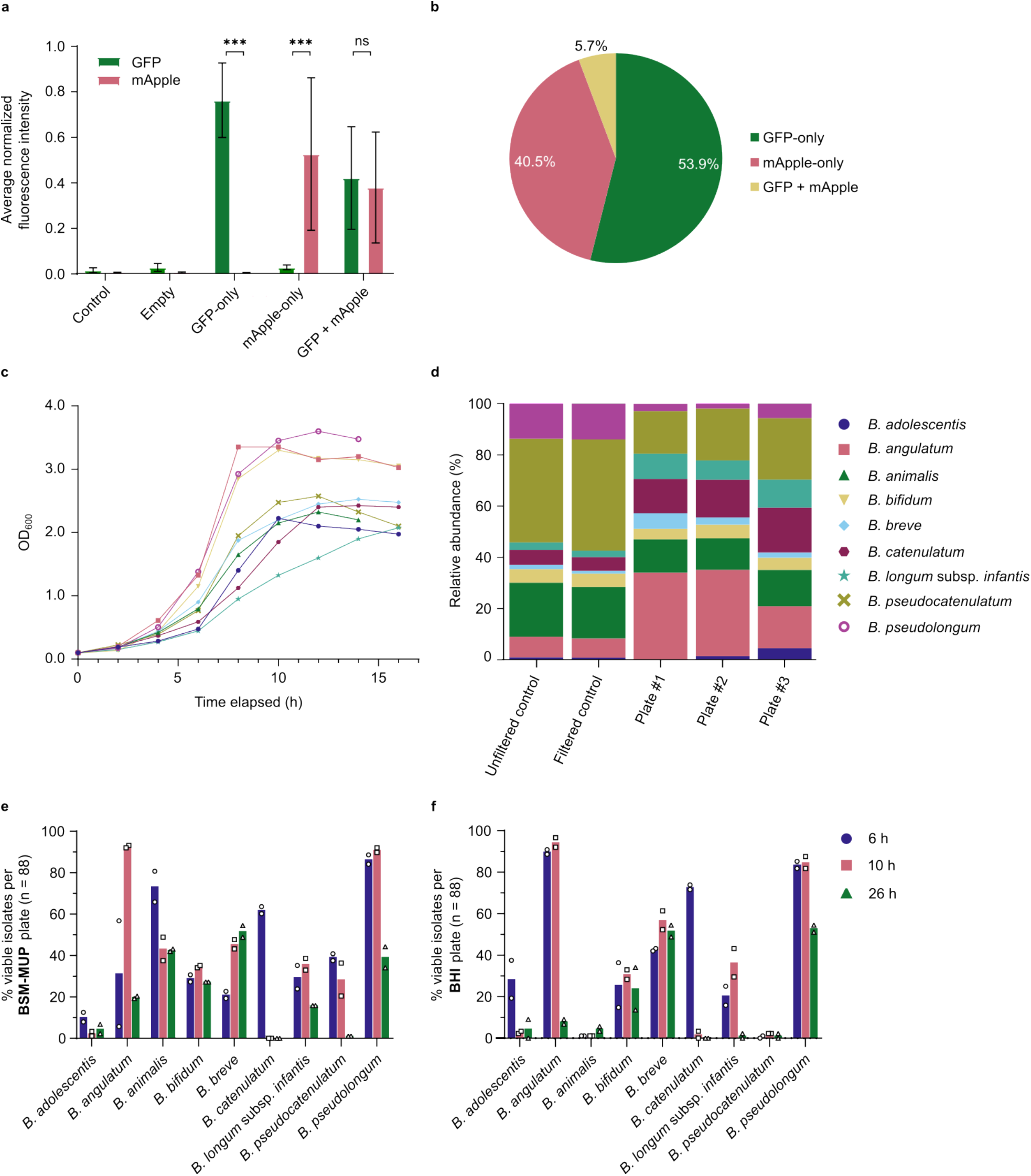
Characterization of the single-cell dispenser’s performance in single-cell isolation and dispensing pure *Bifidobacterium* cultures. (a) Average normalized fluorescence intensity detected from individual wells after dispensing of a mixed GFP-expressing and mApple-expressing *K. pneumoniae* culture. *Control:* non-template control wells. *Empty:* wells with no viable isolates. A Student’s t-test was performed to evaluate the significance of differences in GFP and mApple fluorescence signal intensities (^*\*\*\**,^ *p < 0*.*001*). (b) Proportions of viable isolates dispensed that showed only GFP, only mApple, or both signals. (c) Growth curves of nine *Bifidobacterium* species by monitoring OD_600_ over a 16-hour period. (d) Relative abundances of each *Bifidobacterium* species in the mock community before and after dispensing determined by Nanopore sequencing. *Unfiltered control*: mock community before filtering with 20μm cell strainer. *Filtered control*: mock community after filtering with 20μm cell strainer. (e, f) Percentage of viable isolates per 96-well plate (n = 88) after dispensing pure *Bifidobacterium* cultures at different timepoints into (e) BSM-MUP and (f) BHI.

Next, to investigate whether the single-cell dispenser preserves the global bacterial diversity of samples, we dispensed a mock community of nine *Bifidobacterium* species to test for any biases in its cell-detection algorithm. To obtain the mock community, the *Bifidobacterium* species were cultured separately until their mid-log phase as determined from their growth curves (Fig. 1c) in Bifidus Selective Medium (BSM), a highly selective medium that supports the growth of *Bifidobacterium* while inhibiting other genera such as *Lactobacillus, Streptococcus, Enterococcus* and molds and supplemented with lithium mupirocin, an antibiotic commonly used as a selective agent in *Bifidobacterium* isolation for its high reported specificity (noted “BSM-MUP” here). Notably, three of the species (*B. angulatum, B. pseudolongum* and *B. bifidum*) showed slightly faster growth rates compared to the rest. The nine *Bifidobacterium* species were pooled to form a mock community, then passed through a 20 μm strainer to remove large clumps prior to dispensing into BSM-MUP. Sequencing of the mock community before and after filtration revealed similar species profiles, indicating that the filtering step did not introduce species-specific bias into the dispensing workflow (Fig. 1d). Although the initial mock community displayed uneven abundances, all nine species were recovered at >1% relative abundance in post-dispensing plates, including low-abundance members such as *B. adolescentis* (1% in filtered control), demonstrating the single-cell dispenser’s ability to capture all species. Nonetheless, notable shifts in relative abundance were observed following dispensing: *B. angulatum* exhibited a marked increase (+19% on average, *n* = 3), whereas *B. pseudocatenulatum* showed a substantial decrease (−25% on average, *n* = 3) (Supplementary Fig. 2). These differences could be partially explained by cell morphologies, with both *B. animalis* and *B. bifidum* showing more elongated shapes when observed under a microscope (BX23, Olympus) (Supplementary Fig. 3). Additionally, *B. angulatum* was uniquely prone to forming sticky clumps during mid-log phase, which may affect its representation during dispensing. Overall, while single-cell dispensers may not preserve relative abundances, they remain a valuable tool for capturing and isolating the global diversity of complex bacterial communities.

Having assessed its dispensing capabilities, we moved onto evaluating different culturing factors that could affect the dispenser’s isolation rates. All nine *Bifidobacterium* species were cultured separately in BSM-MUP for 6 h, 10 h, and 26 h. At each incubation timepoint, pure cultures were dispensed into duplicate plates containing either BSM-MUP or Brain Heart Infusion (BHI), a general, antibiotic-free medium. For seven out of the nine species, comparable maximum recovery rates were observed for BSM-MUP and BHI (Fig. 1e-f). However, for *B. animalis* and *B. pseudocatenulatum*, maximum recovery rates in BHI were considerably lower (5.7% and 2.3%, respectively) compared to BSM-MUP (65.9% and 40.9%, respectively). These observations suggest that the antibiotics in BSM-MUP did not seem to affect the recovery rates of the different *Bifidobacterium* species. When comparing between different incubation periods, *Bifidobacterium* species show a maximum recovery rate corresponding to different growth phases. Considering only the BSM-MUP plates, some species such as *B. adolescentis, B. animalis* and *B. catenulatum* were maximally isolated in their early log phase (6 h, Fig. 1e) while others were better recovered in their mid-to late log phase at 10 h. At 26 h, the recovery rates were mostly lower for all the species, except for *B. breve*. Given such divergence observed, we determined that varying the incubation period would be an essential component in isolating *Bifidobacterium* for higher recovery rates and species-level resolution.

### Successful isolation of diverse *Bifidobacterium* from enriched fecal samples using single-cell dispensing

Before proceeding with fecal sample dispensing, we first checked whether BSM-MUP, despite its ability to grow pure cultures of *Bifidobacterium* species (Fig. 1e), is able to effectively enrich for *Bifidobacterium* in fecal samples compared to alternatives. We compared BSM-MUP with the base medium, BSM, and BSM added with the manufacturer’s supplement (BSM-SUP). The BSM supplement is a mixture of three different antibiotics, including polymyxin B, that inhibits a range of non-*Bifidobacterium* bacterial genera such as *Bacilli, Enterobacteriaceae*, and *Pseudomonas*. A human stool sample (SPMP #1) was filtered and diluted 10^4^ times in each medium and incubated for 21 h (37°C, anaerobic conditions). Each culture was then dispensed separately using the single-cell dispenser into the corresponding media. From 72-hour post-dispensing observations, BSM and BSM-SUP showed similar recovery rates (36.4% and 40.9%, respectively), while BSM-MUP yielded the lowest average number of viable isolates (19.7%) (Fig. 2a). To verify that the selective media enriched specifically for *Bifidobacterium* species, a random sample of isolates (n = 10) from a single plate of each medium were subjected to 16S Sanger sequencing. Surprisingly, despite showing higher recovery rates, the BSM and BSM-SUP isolates sampled were all *Enterococcus* (Supplementary Table 2). In contrast, the BSM-MUP isolates selected all belonged to the *Bifidobacterium* genus, confirming that BSM-MUP is more suitable than BSM and BSM-SUP in enriching for *Bifidobacterium* species from fecal samples prior to dispensing.

**FIG 2.**
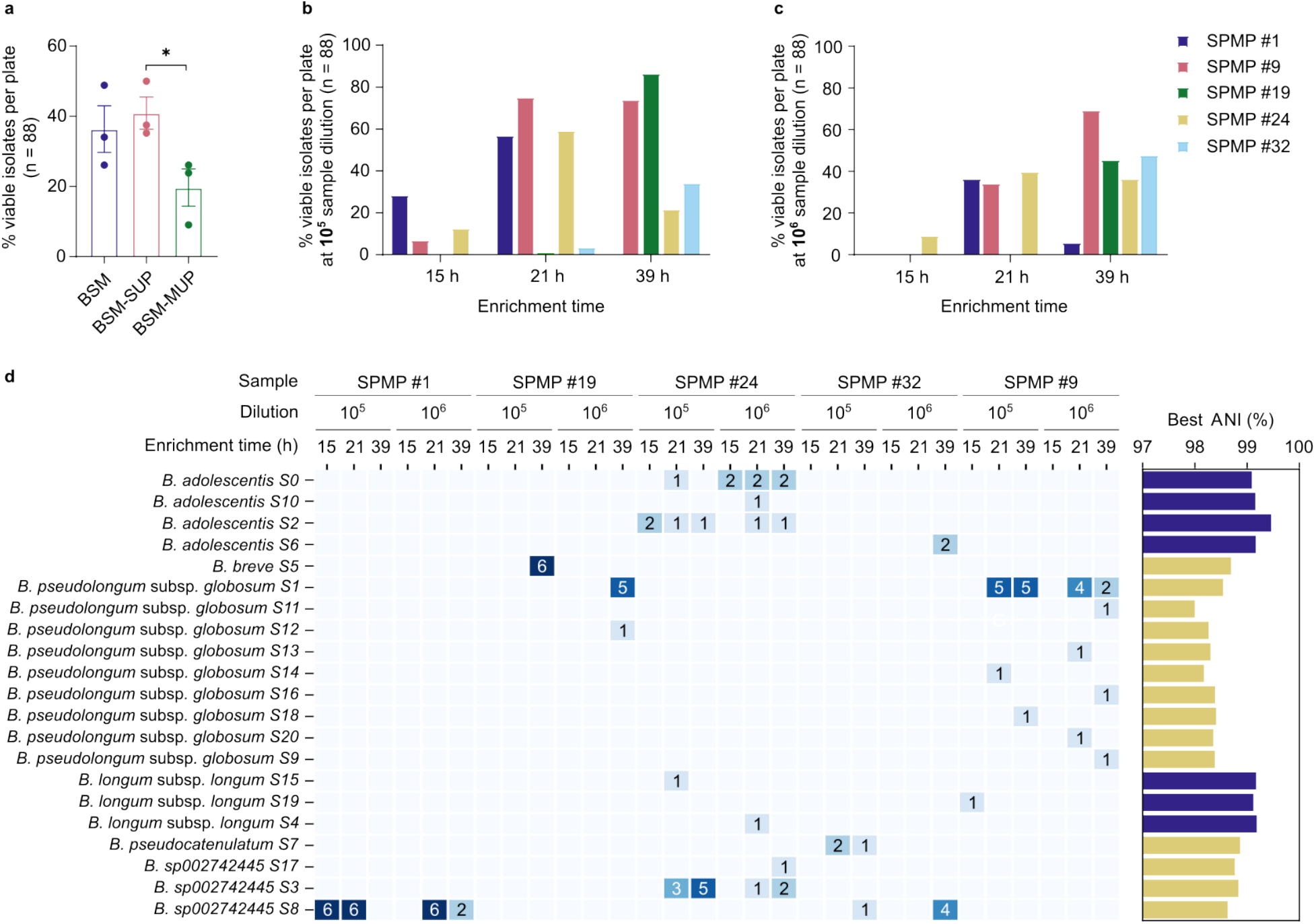
Isolation of *Bifidobacterium* from enriched fecal samples. (a) Percentage of viable isolates per 96-well plate (n = 88) after dispensing a 1:10^4^ stool sample enriched in BSM, BSM-SUP and BSM-MUP for 20 h. Student’s t-test was performed to evaluate the significance of differences recovery rates of each enrichment medium (^*\**^, *p < 0*.*05*) (b, c) Percentage of viable isolates per 96-well plate (n = 88) after dispensing five (b) 1:10^5^ and (c) 1:10^6^ SPMP samples enriched in BSM-MUP for 15 h, 21 h and 39 h. (d) (left) Lineage level-clustering (ANI = 99.9%) of 96 sampled isolates grouped by sample ID, dilution factor, and enrichment time. Each row represents a lineage and each column represents a dispensing run (sample/dilution/enrichment time). Heatmap annotations display the number of isolates obtained. (right) Barplot shows the best ANI obtained for each lineage against all available *Bifidobacterium* genomes within the GTDB R220 database. *Blue*: ANI > 99.0%. *Yellow*: ANI < 99.0%.

Moving forward with BSM-MUP as the chosen enrichment medium, we attempted to obtain isolates from five fecal samples in the *Singapore Platinum Metagenomic Project (SPMP)* cohort (35). Based on our prior observation that different *Bifidobacterium* species show variable growth capability, we tested two different dilutions of the initial samples (10^5^ and 10^6^), and three enrichment times before dispensing (15 h, 21 h and 39 h). In total, 622 isolates were obtained across SPMP samples, enrichment times and dilutions (Fig. 2b-c, Supplementary Table 3). At 15 h of enrichment, only a small number of isolates were obtained for SPMP #1 (10^5^, 28.4%), SPMP #9 (10^5^, 6.8%) and SPMP #24 (10^5^, 12.5%; 10^6^, 9.1%), suggesting this time point might have been too early for *Bifidobacterium* to proliferate. Interestingly, enrichment at 21 h yielded the peak recovery rate for samples SPMP #1 (10^5^, 56.8%), SPMP #9 (10^5^, 75.0%) and SPMP #24 (10^5^, 59.1%) while 39-hour enrichment recovered the most isolates from samples SPMP #19 (10^5^, 86.4%) and SPMP #32 (10^6^, 47.7%). The differing recovery profiles at the various enrichment times examined suggest the divergence in *Bifidobacterium* profiles of each enriched sample, which will further be supported by whole-genome sequencing analyses of the isolates. The identity of 151 randomly sampled isolates was verified using PCR amplification of the *xfp* gene, a gene exclusively found in *Bifidobacterium* species (Supplementary Fig. 4). A vast majority (149/151, or 98.7%) of the isolates showed a PCR product at the expected 235-bp band for *xfp*, suggesting their identity in the *Bifidobacterium* genus.

DNA extraction, whole-genome sequencing and genome assembly were performed on 96 out of the 622 isolates. Six and two isolates were picked per plate for plates showing more and less than 20% recovery rates, respectively. All 96 isolates were barcoded individually and sequenced on a single Oxford Nanopore MinION flow cell. For each isolate, genomes were assembled and classified at the species level against reference genomes in the GTDB R220 database (36). Across these isolates, a total of six *Bifidobacterium* species and subspecies were found, including *B. adolescentis* (n = 16), *B. longum* subsp. *longum* (n = 4), *B. breve* (n = 6), *B. pseudolongum* subsp. *globosum* (n = 29), *B. pseudocatenulatum* (n = 3) and *Bifidobacterium sp002742445* (n = 37). Leveraging our high-quality genomes, we clustered the isolates using pairwise local alignments identity (Supplementary Fig. 5), identifying 21 unique lineages (ANI = 99.9%) among which the majority (n = 14) have an ANI of less than 99% against publicly available genomes, confirming the significant novelty within Asian microbiomes (Fig. 2d). A total of eight different lineages of *B. pseudolongum* subsp. *globosum* (*B. globosum*) were yielded at different enrichment times (21 h and 39 h) and dilutions in sample SPMP #9. Similarly, three distinct *B. adolescentis* lineages were isolated from sample SPMP #24 across culturing conditions, displaying the possibility to obtain not only diverse species, but also intra-subject species diversity with our isolation framework, a component still hard to explore with metagenomic data. Different species also appeared at varying sample dilutions. At 39 h, only *B. breve* was isolated for the 1:10^5^ sample, while the same enrichment time yielded only *B. globosum* for the 1:10^6^ sample; indicating the importance of culturing parameters such as enrichment time and dilution factor when utilising a single-cell dispenser in a culturomics workflow.

Besides the 14 lineages with an ANI of less than 99%, we wanted to determine whether the other lineages were matched to metagenomic sequences or whole-genome sequences from isolates. Interestingly, 6 out of 7 of the *B. adolescentis* and *B. longum* subsp. *longum* isolates were novel when compared to only isolate-derived sequences in the GTDB database (Supplementary Fig. 6). We next compared our isolate assemblies with *Bifidobacterium* MAGs produced from the same stool samples using hybrid assemblies (35). Out of the 11 isolate species/sample pairs, eight pairs were exclusively found within our current dataset, three were found in both, and three were found only within the metagenomic dataset. These results confirm the importance of culturomics in the obtention of a comprehensive view of the species genomic diversity within microbiomes.

## DISCUSSION

### Performance of single-cell dispenser

This study has successfully characterized some key capabilities of a microbial single-cell dispensing system, including its doublet frequency and dispensing bias. To determine the doublet frequency, a dual fluorescence detection method was adopted by dispensing two fluorescent *K. pneumoniae* strains simultaneously, resulting in a doublet frequency estimate of 11.5% (Fig. 1b). The doublet frequency observed using this method was higher than by visual inspection of the nozzle images post-dispensing, suggesting that the latter could be underestimating the true number of droplets. This observation is likely due to the low camera resolution and dark edges of the cartridge obscuring cells in the ROI (Supplementary Fig. 7), as well as the possibility of overlapping cells recognized as a “single cell”. Other methods were also tested, including observation of double colonies when cells were dispensed onto agar-filled wells and identification via growth curves, but neither was useful in estimating the doublet frequency (Supplementary Fig. 8). A different approach was previously described where GFP-transformed *E. coli* cells were dispensed onto glass slides and counted using a fluorescence microscope (26). The method estimated a doublet frequency of 4.0 ± 0.4%, which is lower than the frequency observed in this study. This lower frequency could be explained by the fact that cells may be dispensed into the same spot, leading to the possibility of multiple cells contributing to the same fluorescent spot on the glass slide. The dual fluorescence detection method used in this study removed such biases from affecting the doublet frequency estimate. Through this study, we also show that cartridges could be reused by washing them three times or more with PBS, resulting in relatively low false positive rates for the next sample (Supplementary Fig. 1). In addition, each PBS wash only took approximately three minutes to complete, making it relatively time-efficient. However, the upper limit of how many times the cartridge can be washed with PBS remains to be further tested, as leakage was observed after several washes.

We have also shown in this study that the single-cell dispensing system could preserve the global diversity of a mixed bacterial community. While substantial disparities were observable in our sample before dispensing, all nine species were obtained at relatively high abundance (>1%) on all triplicates, indicating our ability to capture the comprehensive *Bifidobacterium* species diversity with camera-based automatic dispensers (Fig. 1d). However, the single-cell dispenser was found to be unable to maintain the relative abundances of all *Bifidobacterium* species in the mock community (Supplementary Fig. 3), that could be linked to the dispenser cell detection system due to unequal cell shapes or cells-cells interactions such as clumps formation. For instance, as observed under a microscope, two of the four species that showed a decrease in relative abundance, *B. animalis* and *B. bifidum*, have a more elongated shape compared to the rest of the species (Supplementary Fig. 2). To test this potential bias, further experiments that vary the cell-detection parameters of the platform, such as roundness, would be needed. Moreover, other sources of bias, such as species-specific oxygen sensitivity, cell size and plasma granularity, could be further tested to provide a more thorough understanding of capabilities of the single-cell dispensers.

### High-throughput single-cell isolation of *Bifidobacterium*

Having shown that the single-cell dispenser is capable of isolating diverse *Bifidobacterium* species, we aimed to examine different culturing factors that could affect its recovery rates. By dispensing pure cultures of the different *Bifidobacterium* species into both BSM-MUP and BHI, we were able to compare the maximum recovery rates between the two post-dispensing media. We observed that for most of the species, the maximum recovery rates were similar between BSM-MUP and BHI, indicating that the impact of the antibiotics in BSM-MUP on isolate survival was minimal (Fig. 1e-f). In addition, for two of the species, *B. animalis* and *B. pseudocatenulatum*, the recovery was better in BSM-MUP compared to BHI, which suggests that BSM-MUP could be more suitable for culturing the isolates post-dispensing. Possible explanations include the fact that BSM is specifically formulated to stimulate the growth of *Bifidobacterium* species (37), while BHI is a general-purpose medium, and that the bacteria were already accustomed to BSM-MUP pre-dispensing. Regardless, we found that BSM-MUP did not affect the recovery rates post-dispensing. In contrast, varying the incubation period pre-dispensing seemed to affect the recovery of individual *Bifidobacterium* species (Fig. 1e). Incubation periods of 6 h and 10 h produced the maximal recovery rates in BSM-MUP for eight of the species, while *B. breve* was best isolated at 26 h. These timepoints correspond to different growth phases of the species (Fig. 1c), which suggests each species can be recovered maximally at a unique growth phase. Thus, we determined that varying the incubation period is essential for obtaining better isolation rates and diversity from a mixed bacterial community using a single-cell dispenser.

To further validate our workflow, we also assessed the performance of BSM-MUP compared to alternatives such as BSM base medium and BSM-SUP. Despite showing a lower recovery rate compared to its alternatives (Fig. 2a), BSM-MUP was the only medium that successfully yielded *Bifidobacterium* isolates at high levels. When random isolates were sampled for each medium, BSM and BSM-SUP isolates were all identified as *Enterococcus* (Supplementary Table 2). Although BSM was previously claimed by the manufacturer to inhibit the *Enterococcus* genus, we suspect that, because BSM was mainly manufactured to allow for quality control of dairy products, it was not able to inhibit the growth of *Enterococcus* species more predominantly found in the gut such as *E. faecium*. On the other hand, mupirocin is a selective agent that has been shown to be effective against *E. faecium* and *E. faecalis* but non-inhibitory to *Bifidobacterium* (38). We also further examined the efficacy of mupirocin by culturing pure *E. faecium* and *Bifidobacterium* species, as well as *E. faecium* isolates identified on BSM-MUP, compared to BSM and BSM-SUP. The results show complete inhibition of *E. faecium* and growth of *Bifidobacterium* species after 48 h and align with the lack of *Enterococcus* and dominance of *Bifidobacterium* in BSM-MUP isolates (Supplementary Fig. 9). In this study, we did not compare our results with TOS-MUP media (trans-galactosylated oligosaccharide propionate, supplemented with mupirocin), also designed for *Bifidobacterium* isolation, but with lower reported specificity compared to BSM-MUP (37). Furthermore, compared to TOS-MUP, which is only available as agar, BSM-MUP allows for the enrichment of fecal samples in liquid cultures. Liquid cultures remove the additional steps of streaking and colony resuspension before dispensing, which makes them more compatible for use with the system. Thus, compared to other media, BSM-MUP shows the best selectivity for *Bifidobacterium* when using fecal samples for high-throughput dispensing.

### *Bifidobacterium* profiles of fecal samples

When the SPMP samples were enriched in BSM-MUP and dispensed using the single-cell dispenser, high recovery rates were observed at different time points, depending on the sample and dilution factor. Peak recovery rates were observed for SPMP #1, #9 and #24 at 21h (10^5^ dilution), while the maximum recovery was at 39h for SPMP #19 (10^5^ dilution) and SPMP #32 (10^6^ dilution) (Fig. 2d). This was likely due to the unique *Bifidobacterium* species profiles of each sample, giving rise to different growth rates and, hence, recovery rates at each time point. Out of the 151 isolates obtained across SPMP samples, dilutions and time points, 149 (98.7%) were shown to be positive for the *Bifidobacterium* marker gene *xfp* (Supplementary Fig. 4). Thus, despite varying recovery rates across time points, the dispensing workflow was able to achieve highly specific isolation of *Bifidobacterium* that can be leveraged for further large-scale isolation studies in the laboratory.

Among the 96 SPMP isolates sequenced, whole-genome analyses revealed a variety of six different *Bifidobacterium* species present across samples, including *B. adolescentis, B. breve, B. longum* subsp. *longum, B. pseudocatenulatum, B. pseudolongum* subsp. *globosum* and *Bifidobacterium sp002742445. B. pseudolongum* subsp. *globosum (B. globosum)* is often considered as a subspecies of *B. pseudolongum*, a species typically found in the bovine rumen (39), even though instances of *B. globosum* have been previously reported in the gut based on a Chinese cohort (40). Some studies have suggested the role of *B. pseudolongum* in reducing triglyceride levels and suppressing non-alcoholic fatty liver disease (NAHLD) complications in murine models (41, 42). Thus, the presence of *B. globosum*, a subspecies of *B. pseudolongum*, in the SPMP cohort provides an avenue for further investigation into the role of these *Bifidobacterium* in Singaporean gut microbiomes. *Bifidobacterium sp002742445* is a candidate species identified in the human gut with high similarities to *B. catenulatum* and *B. kashiwanohense* (43). Despite the recency of its discovery, some studies have suggested beneficial characteristics of the candidate species based on genomic analysis. For instance, *Bifidobacterium sp002742445* was previously found to be the gut bacterial species to have the second highest percentage of polyphenol utilization proteins within the Unified Human Gastrointestinal Genome (UHGG) database, which implies the possible involvement of the species in polyphenol metabolism in the gut (44). In another study, *Bifidobacterium sp002742445* was shown to be among the most enriched species in a Chinese cohort of osteoarthritis patients (45). Hence, future work could further characterize this species, elucidating its roles in the gut of Singaporeans, especially given that it was the most prevalent species isolated from the five SPMP samples.

Across the genomes assembled, a total of 21 strain-level clusters were identified at 0.1% divergence (Fig. 2d). While only a single strain was isolated for some species such as *B. pseudocatenulatum* and *B. breve*, four unique lineages were observed for *B. adolescentis*, with three of the four clusters originating from the same sample (SPMP #24), highlighting intra-subject diversity. Inter-subject diversity was also observed, as three unique lineages were identified for *B. longum* subsp. *longum* and *Bifidobacterium sp002742445* across four samples (SPMP #1, SPMP #9, SPMP #24 and SPMP #32). In contrast, *B. globosum* found in both SPMP #9 and SPMP #19 also belonged to the same strain (*B. globosum* S1). As such, both cross-sample similarity and diversity were observed between the isolates. Additionally, among the 21 lineages isolated, 14 showed low similarity to publicly available genomes in the GTDB database (ANI < 99%, Fig. 2d). These potentially novel strains were found across four different species, including *B. breve, B. globosum, B. pseudocatenulatum* and *Bifidobacterium sp002742445*. Further analyses could be done to characterize these genetically distinct isolates. Taken together, these findings validate the capabilities of using a dispenser-assisted workflow in capturing both the species and strain-level diversity of *Bifidobacterium species* in fecal samples.

Lastly, by examining the species profiles across different sample dilutions and enrichment periods, the importance of these pre-dispensing culturing parameters in successful isolation of diverse *Bifidobacterium* species was highlighted. For example, the different enrichment periods (15h, 21h and 39h) yielded different species profiles in the 1:10^5^ SPMP #24 sample (Fig. 2d). This reconfirms the importance of varying the pre-dispensing incubation periods to obtain more species diversity, as the *Bifidobacterium* species showed varied growth rates (Fig. 1c) and recovery across growth phases (Fig. 1e). Similarly, the dilution factor when preparing each sample may also play a role in recovering different species of *Bifidobacterium*. By comparing between the 10^5^ and 10^6^ dilutions for sample SPMP #19, it was observed that the two dilutions yielded completely different species (Fig. 2d). Further studies could experiment with other dilution factors and enrichment periods to provide a better understanding of the influence of these parameters on the variety of isolates.

## CONCLUSION

Our study has highlighted the utility of a single-cell dispensing technology to accelerate the isolation of a genus of interest from environmental samples. The performance of the single-cell sorter was first evaluated, revealing a low doublet dispensing rate. Besides, BSM-MUP was demonstrated to effectively enrich fecal samples for *Bifidobacterium*, with the enrichment duration playing a key role in recovery rate of the single-cell dispenser. Building on these findings, we successfully isolated 622 viable strains from fecal samples, all of which distinct species and unique strains of *Bifidobacterium* were discovered. Future work will be required to further optimize the current isolation workflow using a larger number of fecal samples. Moreover, the presence of unique species such as *B. globosum* and *Bifidobacterium sp002742445* as well as potential novel *Bifidobacterium* strains in the SPMP samples prompt further characterization of these *Bifidobacterium* to uncover the roles they play in the gut microbiome. In this context, the systematic and accurate genomic characterization enabled by this isolation workflow will provide the required dataset to obtain a population-level description of corresponding species and strains. While case-specific modifications will likely be required for the isolation of other species, we hope that the protocol established here can serve as a template that can be leveraged for the high-throughput isolation of other bacterial genera, paving the way for a more precise characterization of the genomic diversity within different microbiomes.

## MATERIALS AND METHODS

### Bacterial strains and growth conditions

Two strains of *Klebsiella pneumoniae* SGH10 were obtained as a gift from Prof. Gan’s lab to be used for determining the doublet dispensing frequency with the single-cell dispensing system. Both *K. pneumoniae* strains were transformed with pKPC2 (GenBank, MN542377), each with different fluorescence genes (sfGFP1 or mApple). These strains were cultured separately in Luria Bertani (LB) broth with 50 µg/ml kanamycin at 37°C for 72h in an aerobic condition. In addition, a total of nine bacterial strains of *Bifidobacterium* species from DSMZ were also used: *B. catenulatum* subsp. *catenulatum* B669 (DSM 16992), *B. adolescentis* E194a (Variant a; DSM 20083), *B. angulatum* B677 (DSM 20098), *B. pseudolongum* PNC-2-9G (Biotype a; DSM 20099), *B. animalis* subsp. *animalis* R101-8 (DSM 20104), *B. breve* S1 (DSM 20213), *B. longum* subsp. *infantis* S12 (DSM 20088), *B. pseudocatenulatum* B1279 (DSM 20438), and *B. bifidum* Ti (DSM 20456). All *Bifidobacterium* species were maintained on Bifidus Selective Medium supplemented with lithium mupirocin (BSM-MUP) (Millipore, Darmstadt, Germany) added with 0.01% L-cysteine (Sigma-Aldrich, Darmstadt, Germany) under anaerobic conditions (N_2_ (75%), CO_2_ (20%), and H_2_ (5%)) at 37°C.

### Growth curves and OD_600_-CFU relationships

To assess the doublet dispensing frequency of the single-cell dispenser, the two fluorescent *K. pneumoniae* SGH10 strains were diluted in LB broth to OD_600_ = 0.2 and incubated at 37°C for 9h. At every 1h intervals, OD_600_ was measured to construct a growth curve for each culture, which was then serially diluted (10^−4^–10^−8^) and plated in triplicate spots of 5µl on LB agar for colony-forming unit (CFU) counting. The OD_600_-CFU relationship for each strain was plotted (Supplementary Fig. 10). Approximately 10^7^ CFU/ml of each fluorescent strain was combined to form a co-culture for single-cell dispensing. To assess the single-cell sorter’s dispensing biases, a mock community consisting of the nine *Bifidobacterium* species was used. The OD_600_-CFU relationship of each *Bifidobacterium* species was measured according to the protocol above with slight modification. Briefly, the overnight culture of each species was diluted individually in BSM-MUP broth to OD_600_=0.1 and incubated at 37°C for 16h under anaerobic condition. OD_600_ was then measured at 2h intervals for each species to construct growth curves (Fig. 1c). After performing CFU counting of each species at 6h time point, each species was then re-cultured separately in BSM-MUP broth at 37°C for 6h and ∼3.57×10^8^ CFU/ml of each culture was combined to form the mock community for dispensing.

### Single-cell dispensing

A single-cell dispensing system (B.SIGHT, Cytena GmBH, Germany) was installed within the anaerobic chamber. Samples were kept in a zip-lock bag containing an AnaeroGen 2.5L Sachet (Oxoid, UK) when being transferred between the chambers to reduce the risk of oxygen exposure. All samples were filtered using a 20µm pluriStrainer (pluriSelect Life Science, Germany) before being loaded into the cartridge using non-filtered, sterile 200µl pipette tips. Cell density of each sample was estimated using the number of cells detected by the camera installed in the machine per minute. Samples with excessively high cell density were diluted with pre-reduced PBS. An optimal density of ∼200 to 500 cells/min was used for all fecal samples. Default dispensing settings, including a stroke length of 3µm and a stroke velocity of 75µm/ms, were applied for all dispenses. Cell roundness and size parameters were also set to 0–1 and 0–10 µm respectively for all dispenses. Single cells were dispensed into the first 11 columns (A1–H11) of a Corning round-bottom 96-well plate containing 200µl of liquid media per well. The last column of each plate was used for blanks (wells A12–C12), non-template controls (NTCs; wells D12–G12), and a 400-droplet positive control (well H12). NTC droplets were detected by the camera to be empty and served as a negative control for all dispenses. After dispensing, the 96-well plates were incubated at 37°C.

### Post-dispensing cartridge washing

Upon dispensing, the sample was removed from the cartridge and replaced with 70µL of PBS. The remaining cells inside the cartridge were discharged for ∼3 min by choosing the “Collect samples” function in the x.sight software version 1.4.1 (Cytena GmBH, Germany). The number of leftover cells was detected by the region of interest (ROI) of the live image on the camera and plotted on the “Results” graph. After the three-minute discharge, the PBS was replaced and the cartridge was washed for another 2-4 times until no cell could be detected. A false positive rate was calculated after each washing step by taking the ratio of leftover cells detected per minute to the original cell density detected per minute.

### Fluorescence detection

Six 96-well Corning plates containing fluorescent *K. pneumoniae* dispensed were incubated in aerobic conditions at 37°C for 24h. Fluorescence from individual wells was detected and quantified using a Tecan Spark Multimode plate reader. For sfGFP1 fluorescence, excitation and emission wavelengths were set at 475nm and 515nm, respectively. For mApple fluorescence, excitation and emission wavelengths were set at 560nm and 600nm, respectively. Fluorescence measurements were normalized by the mean background fluorescence of NTC wells, followed by a min-max standardization to account for different fluorescence intensities between sfGFP1 and mApple. Positive fluorescence signal thresholds were defined as two standard deviations from the mean background fluorescence.

### Pure *Bifidobacterium* cell culture dispensing

All nine *Bifidobacterium* isolates mentioned above were individually diluted to OD_600_=0.1 in BSM-MUP broth. Every 6h, 10h, and 26h of incubation, the cell cultures were diluted to 10^−2^ in phosphate buffered saline (PBS) and dispensed into four 96-well plates, with two plates containing BSM-MUP and another two plates containing Brain Heart Infusion (BHI) media (Oxoid, UK). All media and PBS were pre-equilibrated in the anaerobic chamber for at least 48h before dispensing. After 48-72h of incubation, the amount of wells with growth were counted and recorded.

### Stool sample collection

Five independent human fecal samples were obtained from the *Singapore Platinum Metagenomes Project* (SPMP), labelled (in order) SPMP #1, SPMP #9, SPMP #19, SPMP #24 and SPMP #36. Informed consent was obtained from each subject, with 60 Singapore Dollars given as compensation for their participation. All protocols for this study were approved by the National University of Singapore Institutional Review Board (IRB reference number H-17-026) on May 9th, 2017 and renewed until May 31st, 2021. All SPMP samples were collected from healthy subjects using a BioCollectorTM kit (BioCollective, Colorado, USA). Samples were double-bagged and kept in a -20°C polystyrene box. Samples were stored in the anaerobic chamber (N_2_ (75%), CO_2_ (20%), and H_2_ (5%)) before being homogenized, sealed and frozen at -80°C for long-term storage.

### Enrichment of stool sample and dispensing

Each stool sample (100mg) was transferred to a 1.5ml Eppendorf tube and homogenized by pipetting in 1ml of sterile PBS, which had been pre-equilibrated for 48h in the anaerobic chamber. To evaluate the efficiency of different selective media, the homogenized sample (SPMP #1) was then serially diluted in PBS to 10^−3^, and transferred to either BSM, BSM containing supplement (BSM-SUP) (Millipore, Darmstadt, Germany), or BSM-MUP to make 10^−4^ dilution. After 21h of incubation and enrichment, the cell cultures were filtered through a 20µm cell strainer to remove large non-organic residues and serially diluted to 10^−4^ in PBS for dispensing into 96-well plates containing respective media. After 48-72h of incubation, the identity of each species was verified using V1-V9 16S PCR and Sanger sequencing. Once the suitable culturing media and condition have been determined, high-throughput isolation of *Bifidobacterium* species from all stool samples was performed using the above procedure with slight modification. Herein, five SPMP stool samples enriched in BSM-MUP were diluted to 10^−5^ or 10^−6^ and enriched for 15h, 21h and 39h before dispensing into BSM-MUP.

### DNA extraction and purification

To extract DNA from the dispensed samples in 96-well plates, MGISP-960 (MGI Tech Co., Ltd., China), a high-throughput automated sample preparation system, was adopted to automate the extraction process. Post-dispensing liquid cultures in a 96-well Corning plate were transferred to a 1.3ml deep-well plate and centrifuged at 4000 rpm for 15 min at room temperature and pressure. Supernatant was then discarded, leaving cell pellets of size 0.3–0.5 cm. Deep-well plate with cell pellets was sealed and temporarily stored at 4°C. Lysozyme (20 mg/ml) was prepared in a lysis buffer consisting of 1.2% Triton X-100, 20 mM Tris-HCl pH 8.0 and 2 mM EDTA in nuclease-free water (NFW). An in-house MGISP-960 protocol was then run. Briefly, cell pellets were resuspended in a 90 µl lysozyme-containing lysis buffer and incubated at 37°C for 30 min. Both 12.5 µl Proteinase K and 100 µl AL Buffer (Qiagen, USA) were then added for chemical lysis at 56°C for 15 min. After chemical lysis, 170 µl 1× AMPure magnetic beads were added, mixed and incubated on a shaker for 10 min at room temperature. The samples were then moved to a magnetic stand for 5min before all supernatant was removed. The beads were washed twice with 150 µl of 80% ethanol and DNA was eluted in 75 µl nuclease-free water. Post-extraction quantification of the final DNA yield was performed using Qubit™ 4.0.

### Polymerase chain reactions

The components of two polymerase chain reactions (PCR) for the V1-V9 16S region and *xfp* gene fragment are detailed in Supplementary Table 4. The V1-V9 16S rRNA PCR was used to identify species-level taxonomy of the dispensed isolates by Sanger sequencing. The *xfp* gene PCR was used to identify whether the dispensed isolates belonged to the *Bifidobacterium* genus. The following primers were used:

For 16S rRNA PCR (46):

Forward primer: 5’-AGRGTTYGATYMTGGCTCAG-3’

Reverse primer: 5’-CGGYTACCTTGTTACGACTT-3’

For *xfp* PCR (47):

Forward primer: 5’-ATCTTCGGACCBGAYGAGAC-3’

Reverse primer: 5’-CGATVACGTGVACGAAGGAC-3’

The PCR protocols were performed as described in Supplementary Table 5 using a thermal cycler. For the 16S rRNA PCR, PCR products were incubated with 1× AMPure magnetic beads for 5 min and left on a magnetic stand for another 5 min. Solvent was then removed and beads were washed twice with 80% ethanol. 16S rRNA PCR products were then eluted using an elution buffer and run on a Tapestation gDNA screentape (Agilent, USA). The expected band size was 1.5 kbp. PCR products were then sent for Sanger sequencing (1st Base, Singapore) and closest strain-level matches were determined using NCBI Nucleotide Basic Local Alignment Search Tool (blastn) against the 16S ribosomal RNA (Bacteria and Archaea type strains) database (updated: 2025/07/18). For the *xfp-*PCR, PCR products were directly run on a 2% agarose gel supplemented with 1× gel red for visualization. The expected band size was 235 bp.

### Whole-genome Nanopore sequencing

There were two Nanopore sequencing runs in the study. The first run was to compare the composition of the mock community pre- and post-dispensing to observe any genus-specific biases in the performance of single-cell dispenser. For this purpose, DNA samples were prepared following the SQK-NBD114.24 protocol (Oxford Nanopore, UK). Briefly, for each plate with dispensed isolates, equimolar amounts of extracted DNA from each well were pooled together to form a single 400 ng sample. The extracted DNA from the original mock communities (pre- and post-filtering) were also aliquoted as controls. For each sample/control, DNA repair and end-prep were performed by adding Ultra II End-prep enzyme mix and FFPE DNA repair mix as per protocol. The reactions were incubated at 20 °C for 15 min and 65 °C for five minutes. Native barcode ligation was then done by incubating the samples with four native barcodes (NB01-04) and Blunt/TA Ligase Master Mix at room temperature for 20 min. EDTA (10%) was then added to terminate the reaction. DNA was then purified with AMPure XP Beads followed by two 80% ethanol washes and eluted in nuclease-free water after each step. All barcoded DNA was subsequently pooled together for adaptor ligation by incubating with Native Adapter (NA) and Quick T4 DNA Ligase at room temperature for 20 min. The DNA library was then purified once more with AMPure XP Beads followed by two Long Fragment Buffer (LFB) washes and elution in Elution Buffer (EB), as per protocol. DNA yield was determined using Qubit™ 4.0 and 10-20 fmol of the DNA library was loaded onto a Flongle flow cell (Oxford Nanopore, UK) primed with Flow Cell Flush (FCF), 50 mg/ml Bovine Serum Albumin (BSA) and Flow Cell Tether (FCT). Sequencing was run for 24 h. The second Nanopore sequencing run was to identify *Bifidobacterium* species represented in the isolated strains and to construct their genomes. DNA samples were prepared following the SQK-NBD114.96 protocol (Oxford Nanopore, UK). DNA extracted (200 ng) from each of the 96 isolates selected was repaired, end-prepped and barcoded separately (NB01-96), as described previously. DNA was purified with AMPure XP Beads followed by two 80% ethanol washes and eluted in nuclease-free water after each step. The barcoded DNA was then pooled together to form a single sample before adaptor ligation and flow cell loading. For this round of sequencing, a MinION flow cell (Oxford Nanopore, UK) was used and sequencing was run for 72 h.

### Whole-genome analysis

For the first sequencing run, raw sequencing reads were processed and analyzed using Sylph (v.0.6.1) (48) against the GTDB R220 representative genomes database (36). For the second sequencing run, raw sequencing reads were base-called with Dorado (v7.6.7), the latest version of the base caller available at the point of sequencing, generating a total of 9.20Gbp corresponding to an average of 96Mb per sample. For each barcode, reads were processed using an in-house whole-genome assembly pipeline, briefly described as follows. An initial *de novo* assembly was generated with Flye (v.2.9.5) (49), and potential plasmid assemblies were further improved using plassembler (v.1.6.2) (50). Whole assemblies were then reoriented with Dnaapler (v.1.2.0) (51). Resulting assemblies were first screened at the species-level against the GTDB R220 database with GTDB-Tk (v.2.4.0) (52) using the “classify_wf” method. Two genomes with high contamination (> 90%) or low completeness (< 50%) were identified with CheckM2 (v.1.1.0) (53) and removed for subsequent analyses. To obtain precise genomic comparison with all known *Bifidobacterium* genomic diversity, isolate assemblies were compared to all *Bifidobacterium* genomes found within GTDB R220 and themself using Mummer4 (v.4.0.1) (54) nucmer and dnadiff. A threshold of 75% and 50% of minimum genome alignment (on both sides) for self and GTDB comparisons respectively were defined to avoid spurious ANIs. Isolates were clustered using agglomerative clustering with average linkage and a threshold of 0.1% average identity. Comparison with SPMP metagenomic samples was done using MAGs obtained in our prior study. MAGs were compared against the same GTDB R220 database with Skani (v.0.2.2) (55) and assigned a species with only ANI ≥ 95% considered.

## ACKNOWLEDGMENTS

We thank Prof. Gan Yunn Hwen from National University of Singapore for generously sharing her lab’s bacterial strains for this study. This research was supported by the National Research Foundation Investigatorship grant (NRFI09-0015) to Niranjan Nagarajan and the National Medical Research Council Clinical Scientist Individual Research grant (CIRG22jul-0023) to Lye Chien Boon. This work was supported by the A^*^ STAR Computational Resource Centre through the use of its high performance computing facilities.

