## Supplementary Figures for "High-throughput single-cell isolation of *Bifidobacterium* strains from the gut microbiome"

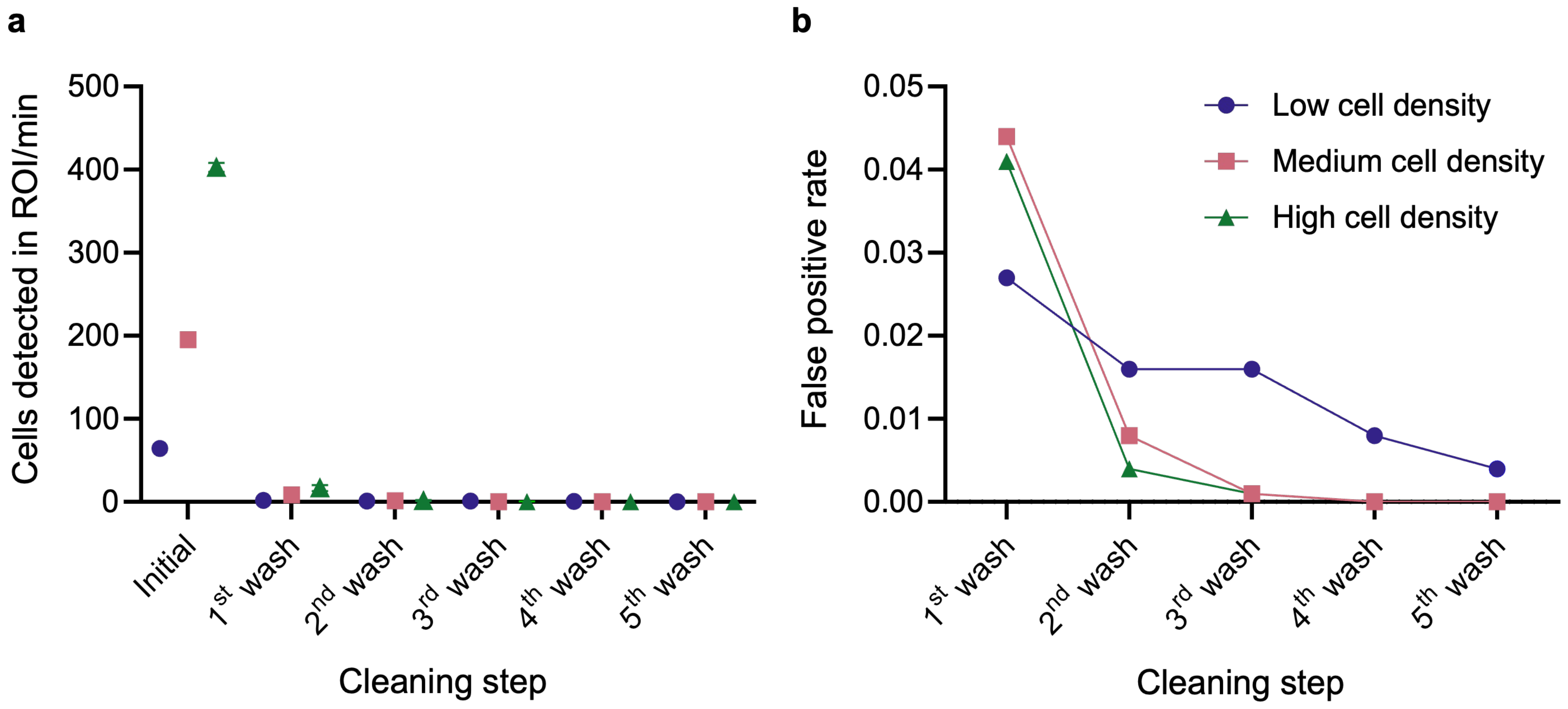


**Supplementary Figure 1. Number of cells detected in the ROI of B.SIGHT camera per minute (a) and false positive rate after multiple rounds of PBS washing (b).** Number of cells detected in the ROI of B.SIGHT camera per minute initially and after each PBS wash were recorded for a single sample diluted to low (blue, 64.4 cells/min), medium (red, 195.4 cells/min) or high cell density (green, mean 402.6 cells/min). Means and standard errors of recordings were reported (n = 3). False positive rate was calculated as the average number of cells detected per minute after each wash divided by average initial count.

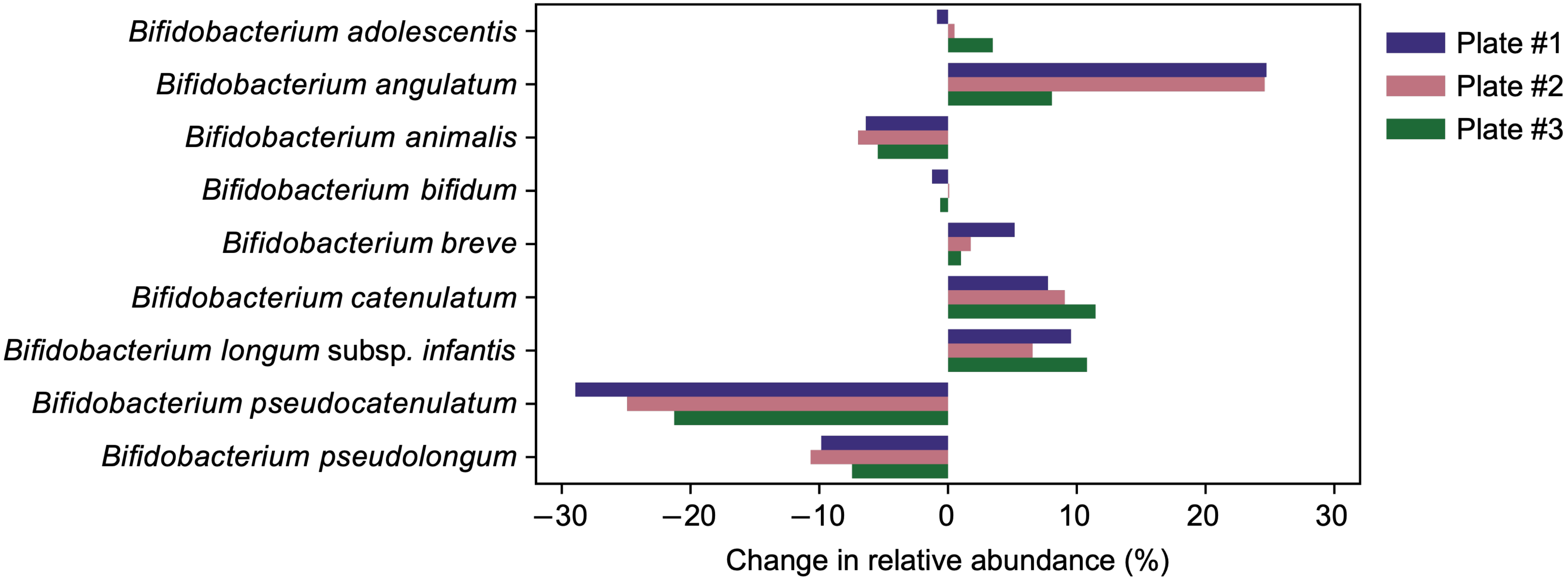


**Supplementary Figure 2. Changes in relative abundances of individual *Bifidobacterium* species in mock community post-dispensing.** Changes were relative to the filtered control mock community.

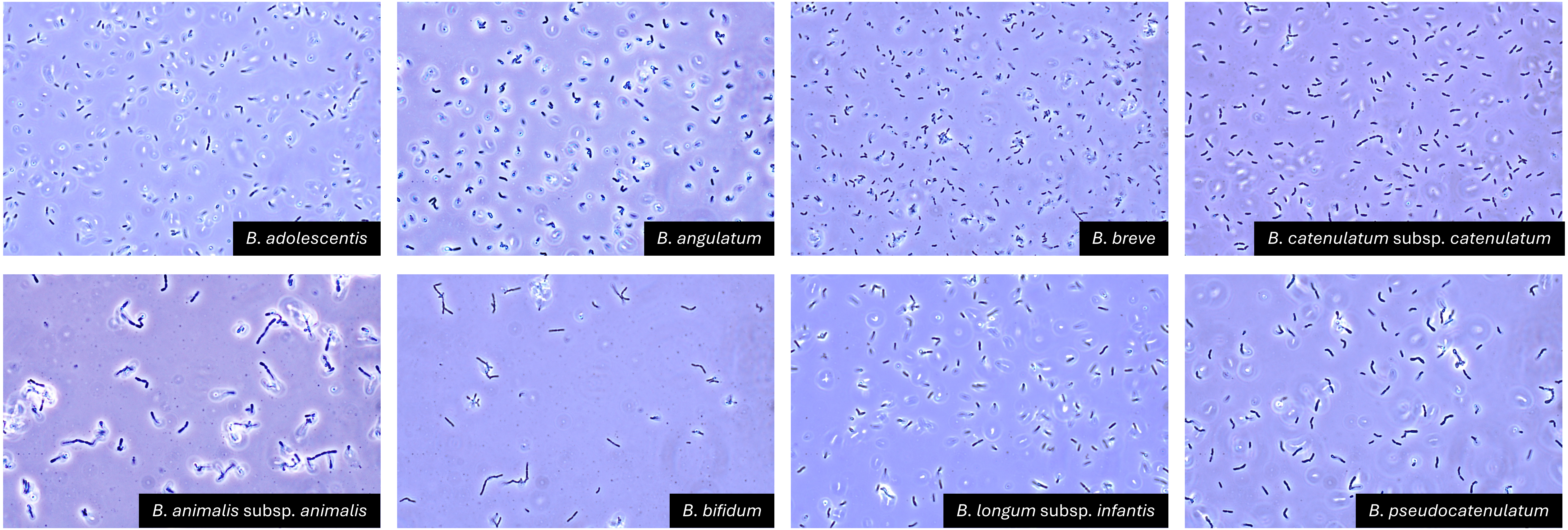


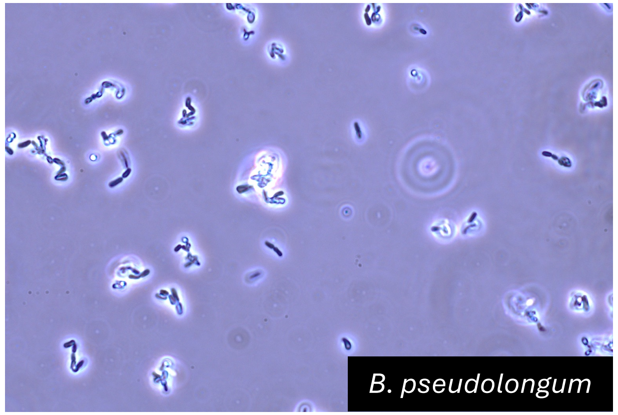


**Supplementary Figure 3. Microscope images of nine *Bifidobacterium* species from the mock community.** Images were taken at the mid-log phase of each species.

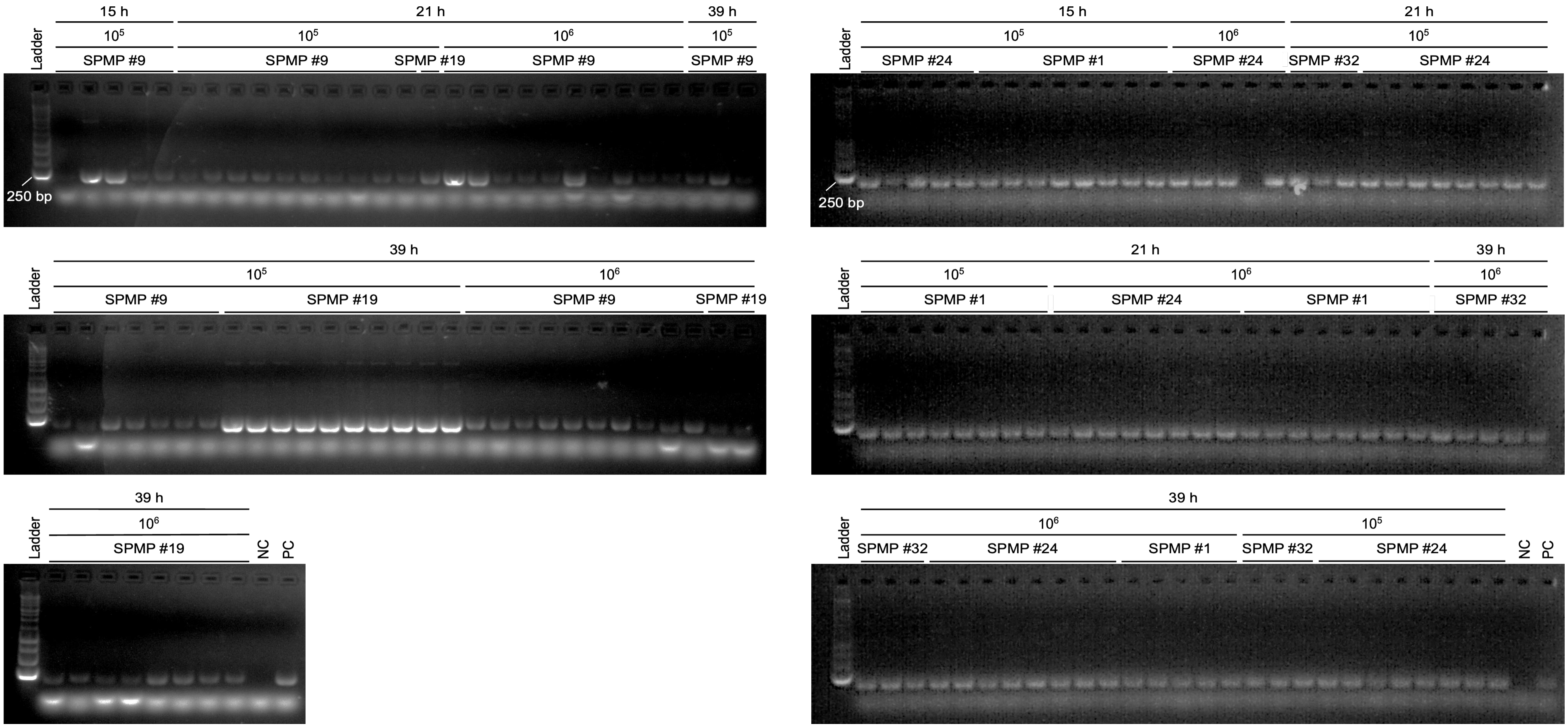


**Supplementary Figure 4. Gel electrophoresis images of *xfp* gene PCR products from SPMP fecal samples.** NC: negative control (NFW). PC: positive control (*B. adolescentis* pure culture).

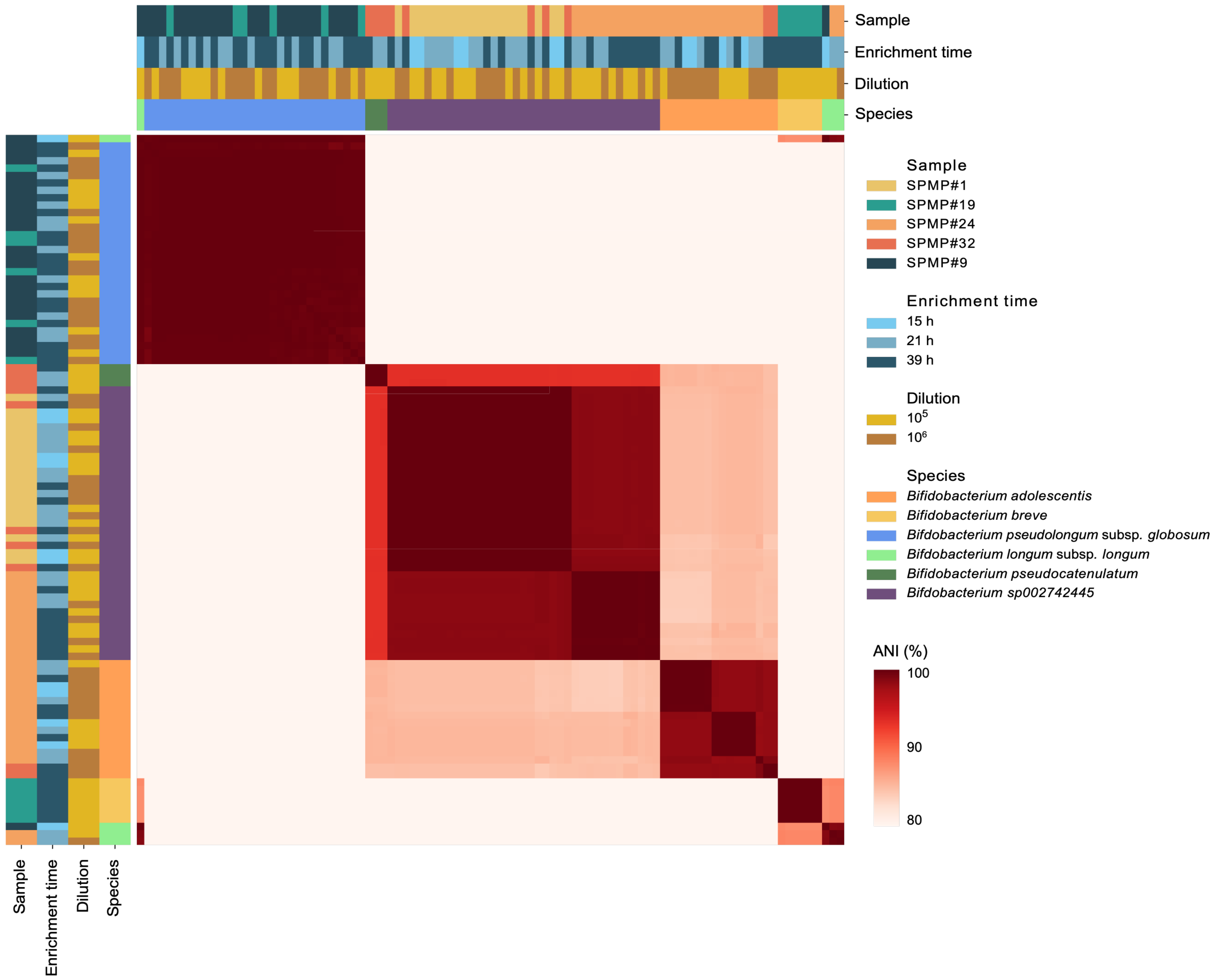


**Supplementary Figure 5. Cluster map showing pairwise ANI comparisons between all 96 SPMP isolates sequenced.** Assembled genomes were compared with each other and ANI was calculated for each pair of genomes. A single isolate (top bar, leftmost) was found to have high ANI with both *B. globosum* and *B. longum* genomes and was further excluded for later analyses.

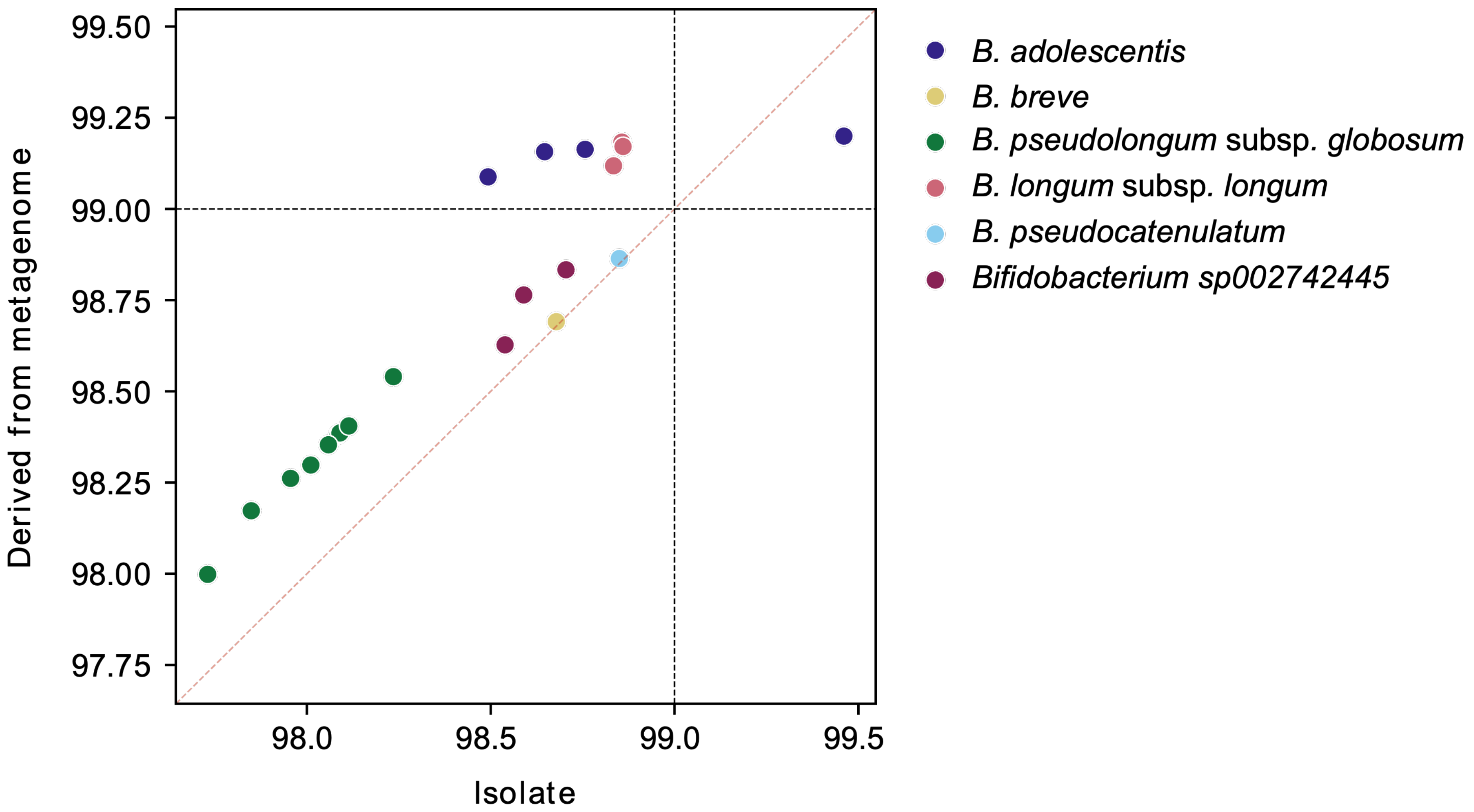


**Supplementary Figure 6. For each dispensed isolate genome, comparison of the best ANI obtained against GTDB genomes with either "Derived from metagenome" or "Isolate" source.** ANI values were obtained from SkANI. Red dotted line represents the diagonal for visual aid. Black dotted lines delimit 99% ANI.

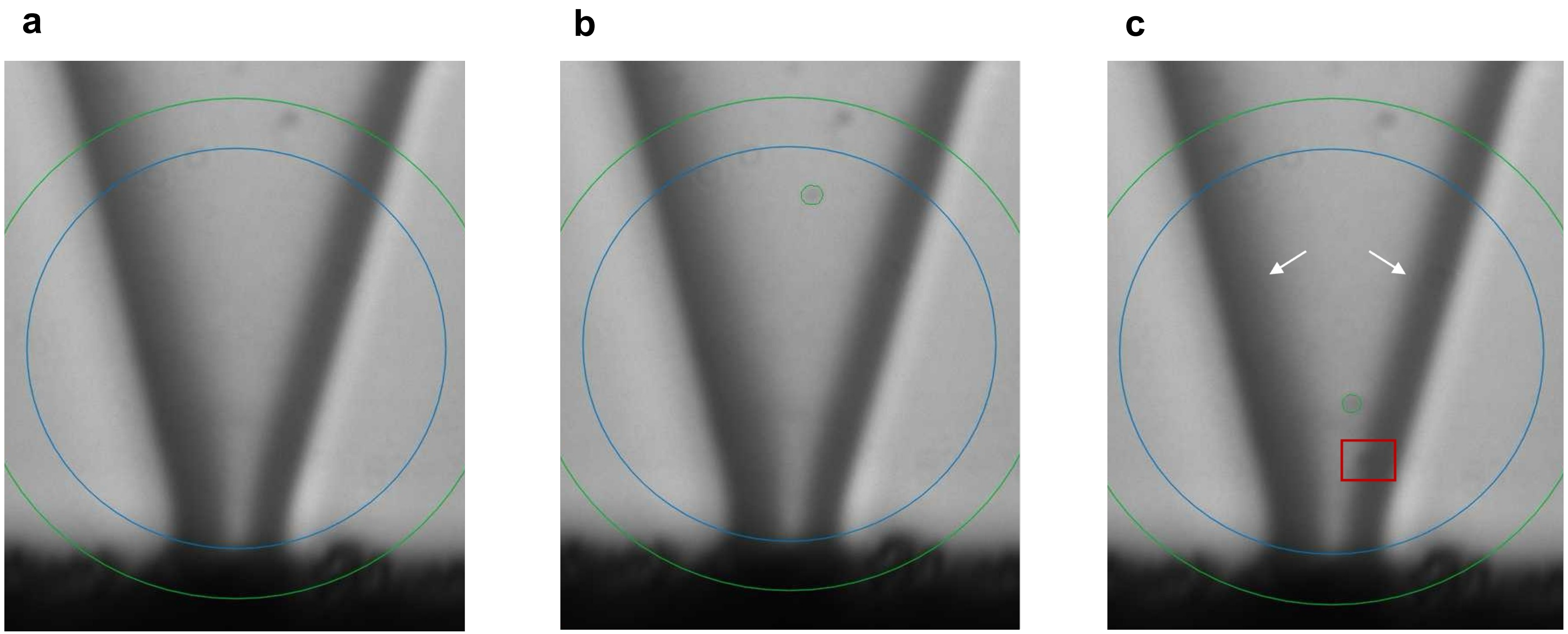


**Supplementary Figure 7. B.SIGHT nozzle images of the same cartridge during the same run.** (a) Empty cartridge. (b) A single cell detected and dispensed (demarcated in green). (c) Two cells (doublets) detected and dispensed (demarcated in green and red). Dark edges of the cartridge (arrows) and low camera resolution could potentially obscure cells in the ROI, leading to droplets with two or more cells.

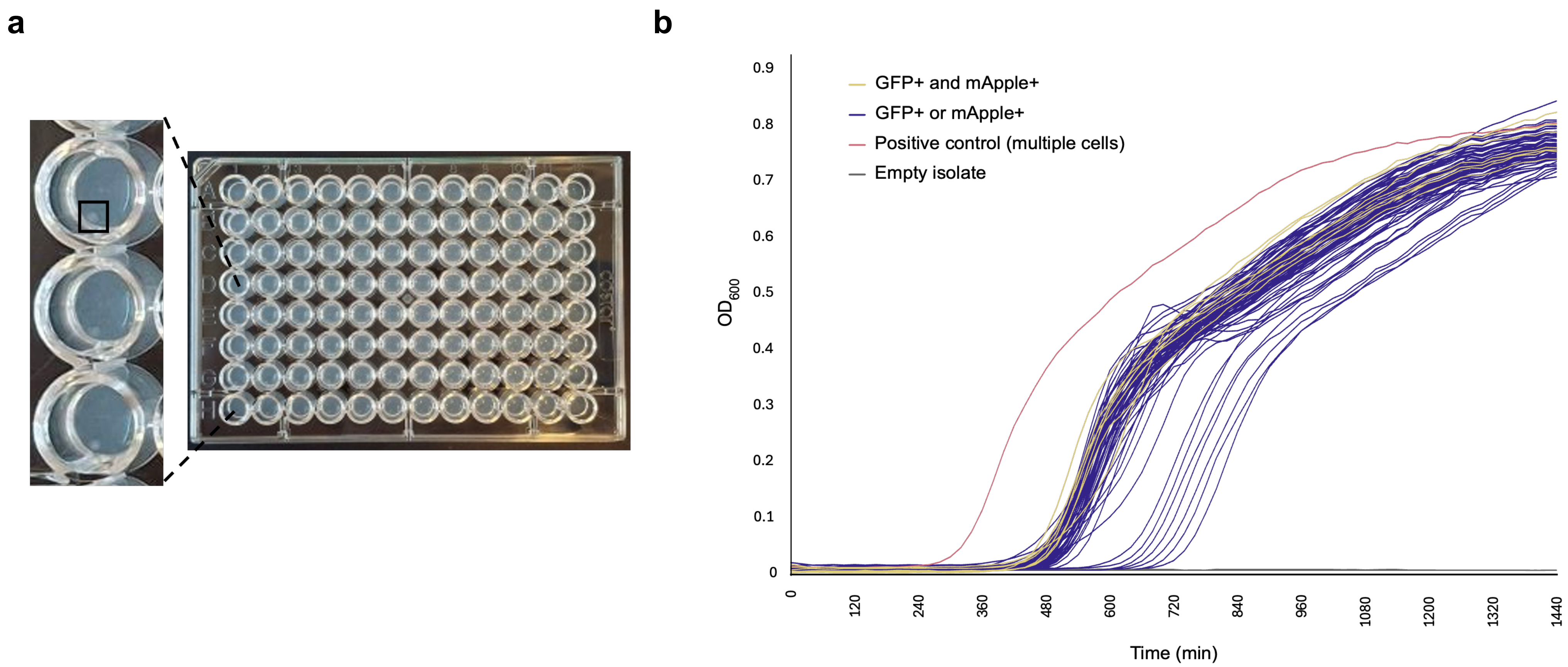


**Supplementary Figure 8. Alternative methods were not useful in detecting doublets.** (a) Visual inspection of colony growth on agar-filled plates. Double colonies could not be visibly discerned when cells were dispensed onto agar-filled wells. All colonies appear as the one shown in the black square. (b) Examination of growth curve variations over 24 hours. GFP+ and mApple+ doublets are indicated in yellow. Isolates with single fluorescence, GFP or mApple, are indicated in blue. Positive control (400 droplets) is indicated in red. Empty isolates with no observable growth are indicated in gray. There was no significant difference between growth curves of doublets (identified using dual fluorescence detection) and single cells.

**a**

| Media | | *E. faecium* | *B. breve* | *B. infantis* | Dispensed *Enterococcus* Isolate | | | | | | |
|---|---|---|---|---|---|---|---|---|---|---|---|
| | | | | | 1 | 2 | 3 | 4 | 5 | 6 | 7 |
| Agar | BSM | + | + | + | + | + | + | + | + | + | + |
| | BSM-SUP | + | + | + | + | + | + | + | + | + | + |
| | BSM-MUP | - | + | + | - | - | - | - | - | - | - |
| Broth | BSM | + | + | + | + | + | + | + | + | + | + |
| | BSM-SUP | + | + | + | + | + | + | + | + | + | + |
| | BSM-MUP | - | + | + | - | - | - | - | - | - | - |

**b**

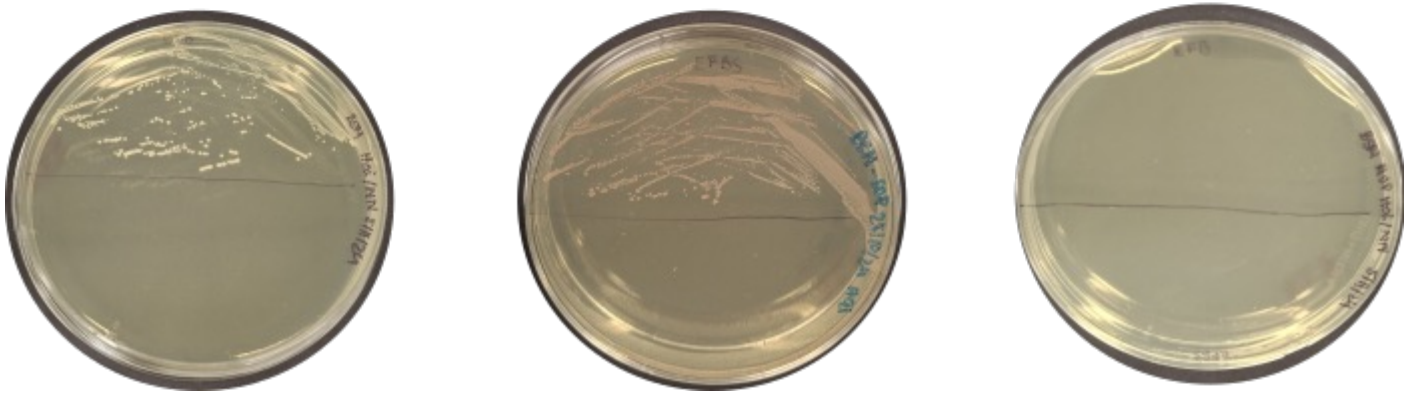

**Supplementary Figure 9. Growth inhibition assay for *Enterococcus* and *Bifidobacterium* species using different BSM-based media.** Isolates were obtained from stool sample and identified using 16S Sanger sequencing. (a) Growth of *E. faecium*, *B. breve* and *B. longum* subsp. *infantis* and dispensed isolates in BSM-MUP compared to BSM and BSM-SUP. "+" indicates growth after 48 h. "-" indicates growth inhibition after 48 h. (b) Images of growth of *E. faecium* streaked on respective agar media. Only the top half of the agar plates were used for streaking.

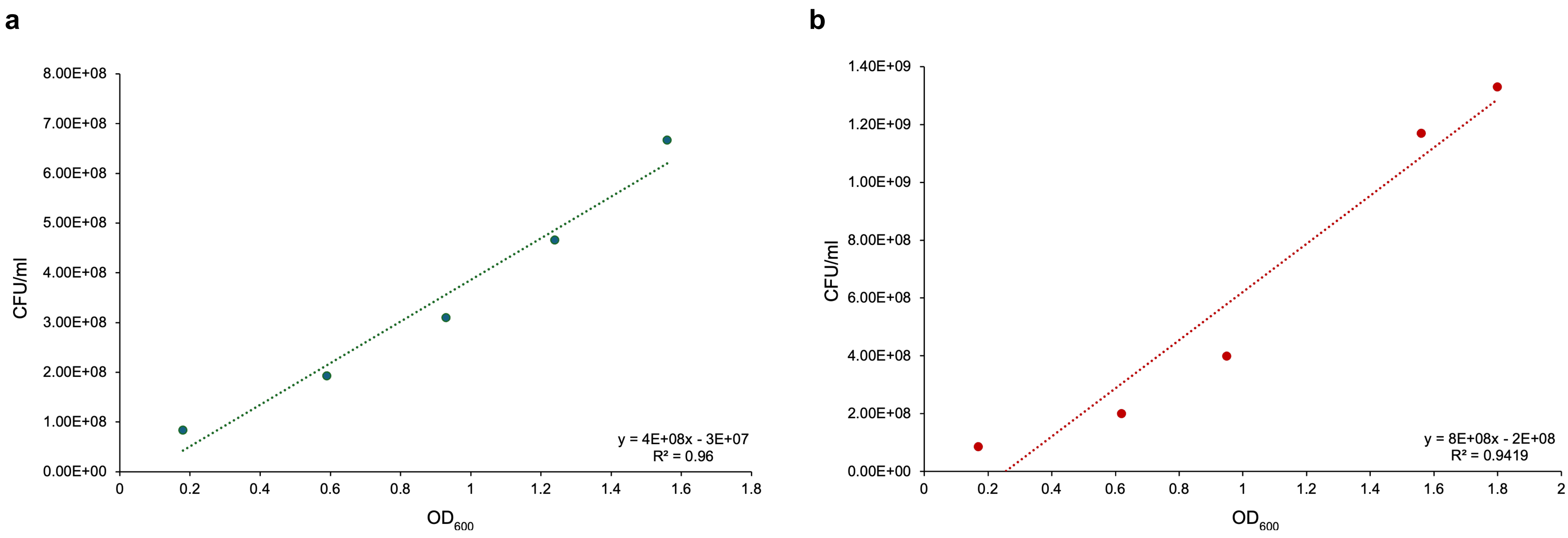


**Supplementary Figure 10. $OD_{600}$-CFU relationship for the two fluorescent *K. pneumoniae* strains.** Each dot represents a $OD_{600}$-CFU paired reading for (a) *K. pneumoniae* transformed with pKPC::sfGFP1 and (b) *K. pneumoniae* transformed with pKPC::mApple. The dotted lines are the best-fit lines, whose equations are show at the bottom right corner of each plot. R-squared values are also given at the bottom right corner of each plot.
